## Supplementary_Information for "Population-level differences in the neural substrates supporting Statistical Learning"

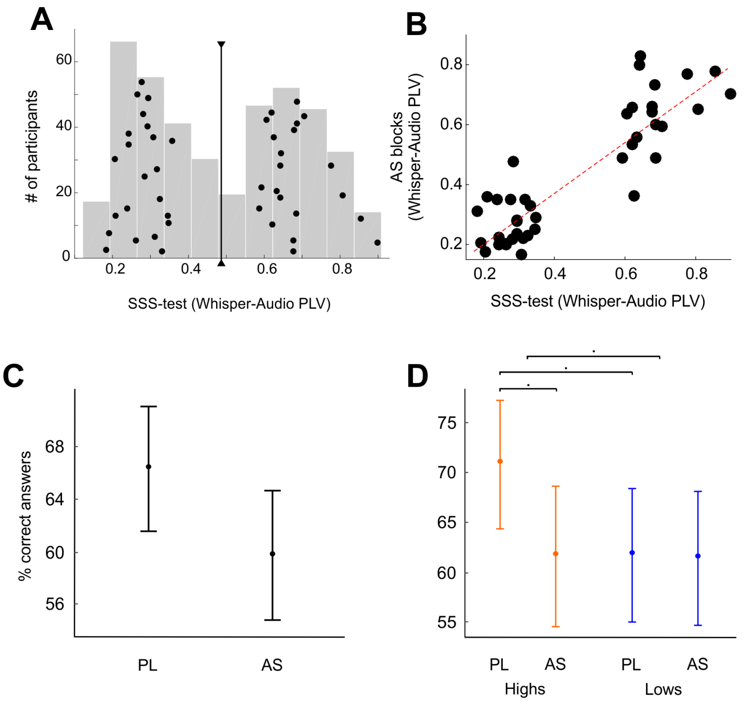


**Fig S1. Behavioral performance in the scanner. (A)** SSS-test outcome. Histogram of 388 PLVs obtained in a previous work[1] with two different versions of the SSS-test. Black dots represent the participants selected to complete the fMRI protocol. Black line represents the threshold value adopted in this work to separate high and low synchronizers: $PLV_{threshold} = 0.49$. A *k-means* clustering algorithm using a squared Euclidean distance metric was applied over this distribution (N = 388). The threshold value is the midpoint between the two clusters' centers. (**B)** Scatterplot displaying participants' PLV during AS inside the scanner as a function of the PLV from the SSS-test. Red line represents the correlation of the data. The correlation is displayed for visualization purposes, to emphasize that the synchronization of low synchronizers is consistently worse than that of highs during the AS block. The correlation within groups remains significant only for high synchronizers ($r_{HIGH}=0.45$ $p_{HIGH}=0.044$; $r_{LOW}=0.21$ $p_{LOW}=0.31$). (**C)** Percentage of correct responses for the statistical word-learning task during PL and AS conditions inside the scanner on the entire sample. (**D)** Percentage of correct responses for the statistical word-learning task during PL and AS conditions inside the scanner for the low (blue color) and the high (orange color) synchronizers. The mixed-model analysis of this dataset yielded a significant difference between conditions (main effect of Condition (PL > AS), $\chi2 = 5.40$, $p < 0.05$)), and a main effect of group close to significance (Highs > Lows; $\chi2 = 3.67$, $p = 0.055$), and a trending Condition*Group interaction ($\chi2 = 2.74$, $p = 0.098$). Dots: model predicted group means. Bars: 95% confidence interval. AS: Articulatory Suppression; PL: Passive Listening.

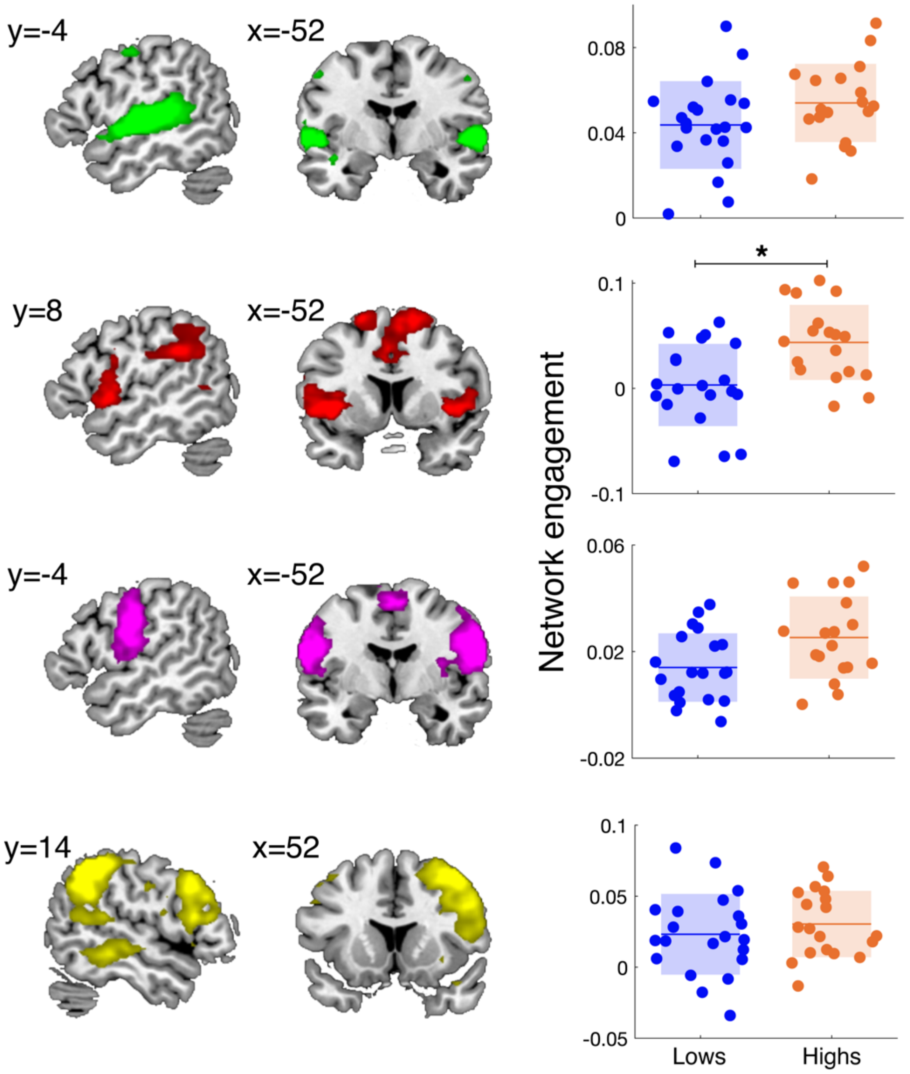


**Fig S2. Brain networks significantly activated during PL.** The different networks are shown over a canonical template with MNI coordinates on the upper portion of each slice. Neurological convention is used with a $p < 0.05$ FWE-corrected threshold at the cluster level and an auxiliary $p < 0.001$ threshold at the voxel level. In addition to the auditory (green) and fronto-parietal (red) networks described in the main manuscript, a sensorimotor (magenta) and a right lateralized fronto-temporo-parietal (yellow) networks were also activated during PL. All these networks were significantly activated during PL for both high and low synchronizers, except the fronto-parietal, which was only active for the high. *$p < 0.05$ Mann-Whitney-Wilcoxon between-group comparison, FDR corrected.

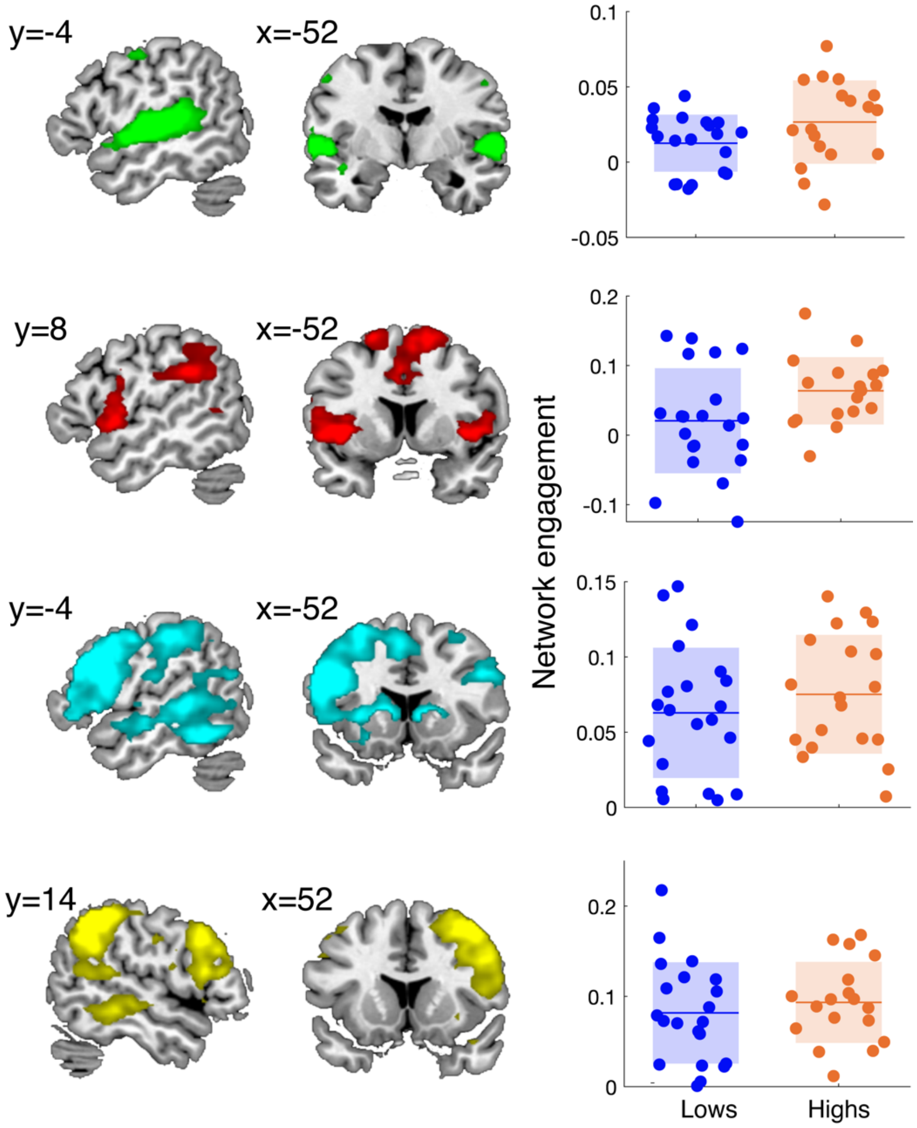


**Fig S3. Brain networks significantly activated during AS.** The different networks are shown over a canonical template with MNI coordinates on the upper portion of each slice. Neurological convention is used with a $p < 0.05$ FWE-corrected threshold at the cluster level and an auxiliary $p < 0.001$ threshold at the voxel level. In addition to the auditory (green) and fronto-parietal (red) networks described in the main manuscript, a left (light blue) and a right (yellow) lateralized fronto-temporo-parietal network were also activated during AS. All networks were significantly activated during AS for both high and low synchronizers, except for the fronto-parietal which was only active for the high and the auditory, which was marginally significant for the lows ($p_{unc} = 0.03$).
